# Serum exosomal miR-150-5p in rheumatic heart disease-associated angiogenesis

**DOI:** 10.1101/2025.06.13.659484

**Authors:** Bhupender Sharma, Shaveta Jain, Lakhwinder Singh, Shruti Sharma, Raman Kaur, Harkant Singh, Seema Chopra, Anuradha Chakraborti

## Abstract

The focus of this study is on Rheumatic heart disease (RHD), an autoimmune consequence of rheumatic fever leading to mitral valve damage. Aim is to explore the role of exosomes derived from RHD patients in promoting angiogenesis during RHD pathogenesis. Exosomes were isolated *via* column chromatography from the pericardial fluid (PF) and serum of RHD patients, pericardial fluid from non-RHD patients undergoing coronary artery bypass grafting (CABG) surgery, and serum of healthy individuals. These exosomes were characterized by transmission electron microscopy (TEM) and western blotting. Human umbilical vein endothelial cells (HUVECs) cultured from umbilical cord(s), were treated with purified exosomes from RHD, CABG patients or healthy individuals. Exosome treatment induced morphological changes in HUVECs, including elongation and tube-like structure formation.

The qRT-PCR gene expression analysis revealed significant upregulation of angiogenic markers (VEGF-A 2.7-fold, MMP2 1.5-fold) and inflammatory cytokine (IL-4 15.17-fold), in HUVECs stimulated with RHD serum exosomes. Conversely, there was downregulation of anti-angiogenic genes (RASA-1 0.65-fold and endostatin 0.4-fold). Notably, elevated levels of miR- 150-5p (34.29 ± 9.524-fold mean difference; p= 0.0007) were detected in serum-derived exosomes from RHD patients compared to healthy controls. Stimulation of HUVECs with RHD serum exosomes resulted in reduction in the expression of TP53 (0.227-fold; p= 0.006), a known target of miR-150-5p. Furthermore, transfection of HUVECs with miR-150-5p mimics induced the formation of angiogenic endothelial tubes, underscoring the role of miR-150-5p in promoting angiogenesis.

**Graphical Abstract:** 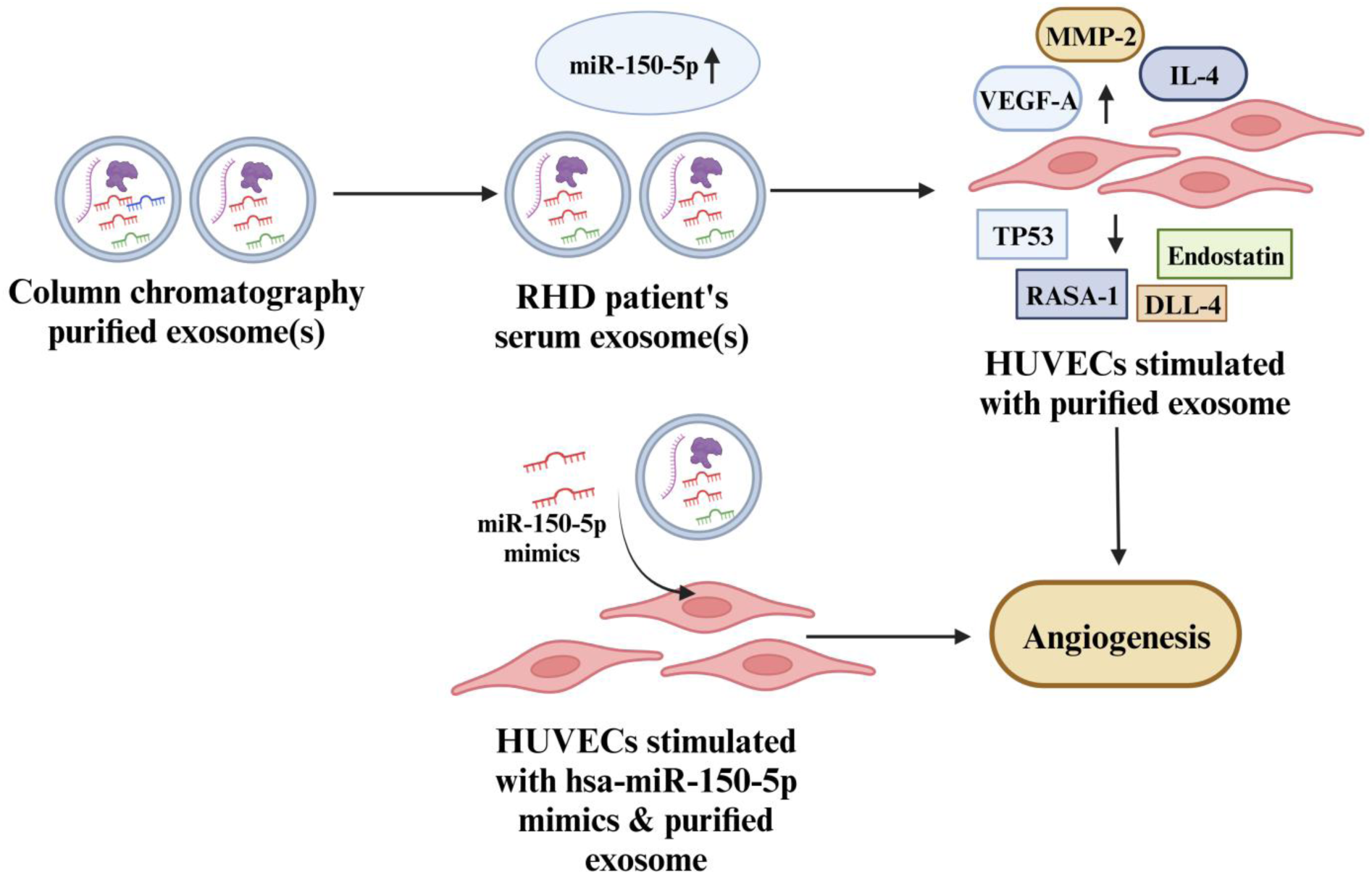

## 1. INTRODUCTION

Rheumatic heart disease (RHD) is an autoimmune disease caused by an infection with Group-A-*Streptococcus* (GAS). The condition primarily affects endothelial cells especially those of mitral valves. An initial GAS infection, commonly causing a sore throat, can develop into acute rheumatic fever (ARF). A single severe episode or recurrent bouts of ARF, may eventually lead to the development of RHD (1). RHD is marked by chronic inflammation of the endocardium and myocardium, resulting in damage to the heart valves (2). A key factor in the progression of RHD is molecular mimicry, a phenomenon in which immune system mistakenly targets the body’s own tissues, like heart valves, as foreign invaders. This happens because antibodies produced by B-cells, which normally target *Streptococcus pyogenes*, also cross-react with human proteins that resemble *Streptococcus pyogenes* antigens (3, 4). The *Streptococcus pyogenes* produces *M*-protein, which share structural similarities with proteins expressed by endothelial cells in the heart and blood vessels. These endothelial cells form a single layer lining throughout cardiovascular system, including heart valves. As a result, antibodies directed against *Streptococcus pyogenes* may also target endothelial cells, leading to inflammation and tissue damage, particularly in the mitral valve (5).

RHD is most commonly seen in children and young adults and is a significant concern in areas with poor sanitation and limited healthcare resources. The absence of specific diagnostic tools and treatment options, aside from valve surgery, has made RHD a significant health burden in many developing nations. The RHD condition is characterized by inflammation, heart valve calcification, angiogenesis, and fibrosis (6, 7). Initially, it was believed that calcification was a result of random calcium buildup on the valve leaflets. However, recent research has shown that calcification in RHD valves is part of an inflammatory process linked to angiogenesis, the formation of new blood vessels (8–10). It has been reported that due to inflammation, endothelial cells are activated to proliferate, migrate, and invade tissue during angiogenesis (11, 12). Angiogenesis, typically occurs during embryonic development, is regulated by growth factors such as VEGF-A, platelet-derived growth factor (PDGF), fibroblast growth factor (FGF) (11, 13), and is involved in various diseases, including RHD (14, 15). The formation of new blood capillaries involves the differentiation of endothelial cells, which form the inner layer of blood vessels (16). In RHD, angiogenesis in heart valves contributes to valve dysfunction (9, 17, 18). Various factors, including exosomes, may influence angiogenesis (19). Exosomes are small, nanoscale vesicles (20-100 nm) released by cells into the extracellular matrix (20) and can accumulate in body fluids (21–28). These vesicles carry bio-molecules like proteins, lipids, and nucleic acids, enabling communication between cells and influencing the surrounding tissue environment. In particular, exosomes help to regulate gene expression and activate signaling pathways in recipient cells (29).

Exosomes are involved in wide range of diseases, including cardiovascular conditions (30–36) autoimmune diseases (33), neurodegenerative disorders, infectious diseases (37), obesity (31) and cancer (38, 39). Our previous research detected α-1-antitrypsin, a protein associated with RHD severity, in exosomes from both serum and pericardial fluid of RHD patients (33). To study exosomes, it is important to isolate and purify them from other cellular debris (40). Among the various molecules carried by exosomes, microRNAs (miRNAs) are of particular interest because they control gene expression by binding to and degrading specific messenger RNAs (mRNAs) (41). The miRNAs are critical in controlling processes like cell growth, differentiation, and apoptosis (42) and angiogenesis (43). Altered levels of circulating miRNAs are associated with a variety of diseases, including cancer (44), neurological disorders (45) and cardiovascular diseases (46, 47). This study investigates the dysregulated miR-150-5p expression in exosomes from RHD patients and explores its potential role in angiogenesis.

## 2. METHODS

### 2.1. Sample collection

The institute ethics committee of PGIMER, Chandigarh, India, approved this study (Approval No: PGI/IEC/2017-729, dated 26.09.2019). Blood and pericardial fluid (PF) were collected from twenty confirmed cases of Rheumatic heart disease (RHD) (10 males, 10 females) undergoing mitral valve replacement surgery at PGIMER’s Cardiovascular and thoracic surgery (CVTS) Department, Chandigarh, India. As a comparison group, PF samples were collected from age and sex matched twenty non-RHD patients (10 males, 10 females) undergoing coronary artery bypass grafting (CABG) surgery at the same department. CABG patients were selected as controls because they do not have an autoimmune condition like RHD. Although during CABG surgery also requires the opening of the chest cavity for grafting, it provides an opportunity to collect PF, which cannot be obtained from healthy individuals under normal circumstances. Blood samples were collected from age and sex matched healthy individuals. Additionally, umbilical cords from five healthy mothers, collected post-delivery at the Obstetrics and gynecology, Department of PGIMER, were used to isolate endothelial cells from human umbilical vein (HUVECs). Samples were collected from 2019-2021 and stored at -80^°^C and study was conducted from 2019-2022. Blood was allowed to clot at room temperature followed by centrifugation (3200 rpm x 15 minutes at 4^°^C) to isolate the serum. Written informed consent was collected prior to sample collections.

### 2.2. Comparison of exosome purification

The pooled serum from RHD patients was diluted with phosphate buffer in a 1:1 (v/v) ration, and then centrifuged at 10,000 rpm for 20 minutes at 4^°^C. DEAE-cellulose (47) and Octyl-sepharose (48) columns were prepared by equilibrating them with their respective starting buffers. The DEAE-cellulose column and Octyl-sepharose column were used for exosome purification, as outlined in our earlier research, with a flow rate of 1.0 mL/min (49–52). Biomolecules bound to DEAE-cellulose column, were eluted using 50 mM potassium phosphate (PP) buffer (pH 8.0) with NaCl (1-10^th^ fractions 0.5 M and 11-20^th^ fractions 1.0 M). Biomolecules bound to the Octyl-sepharose column were fractionated using 50 mM PP buffer (pH 8.0) with a decreasing ammonium sulfate gradient (0.5 M for fractions 1-10), followed by elution with PP buffer (without ammonium sulfate for fractions 11-20). Protein concentrations in the fractions were determined using the Bradford method (53). The eluted fractions were selected based on their absorbance at 450 nm (48). After purification, these fractions were analyzed using transmission electron microscopy (TEM) at PGIMER, Chandigarh, India (TEM 1400 Plus), and AIIMS, New Delhi, India (Morgagni 268D).

### 2.3. Exosomes purification *via* column chromatography

Pooled PF samples from RHD patients were applied to an Octyl-sepharose column using a loading buffer composed of 50 mM PP buffer (pH 8.0) and 1.0 M ammonium sulfate (1.0 M). Thereafter column was rinsed with the same buffer to eliminate any unbound or weakly bound proteins. Exosomes that were retained on the column matrix were then eluted with 50 mM PP buffer (pH 8.0). The eluted and purified exosome fractions were pooled and desalted using centrifugal filters with a 10 kDa molecular weight cut-off (MWCO) (Amicon ultra, Merck Millipore). Concentrated exosome-containing fractions (1 mL, 0.6 mg) were then loaded onto a Sephadex G-100 column (Pharmacia fine chemicals) for size-based fractionation. After performing Sephadex G-100 column chromatography, the fractions (25 × 2 mL) were analyzed for protein content at A280 nm and acetylcholinesterase activity. This column chromatography method was similarly applied to purify exosomes from RHD patients (serum and PF samples CABG patients (PF samples) and healthy individuals (serum samples). The purified column fractions were stored at -20^°^C for future use.

#### 2.3.1. Acetylcholinestrase assay

To test acetylcholinesterase activity, 20 µL of the purified exosome samples were added to a 96-well plate containing 1.25 mM acetylthiocholine and 0.1 mM 5,5-dithiobis-(2- nitrobenzoic acid) (DTNB), with a final volume to 150 µL. The mixture was incubated at 37^°^C for 30 minutes to allow the reaction to occur and absorbance at 450 nm were recorded using a multi-mode reader (Infinite pro 200 Tecan) (48, 49, 52) to calculate enzyme activity using a standard formula.

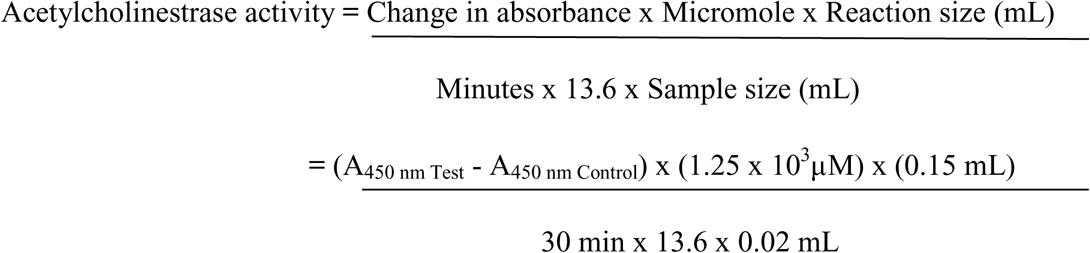

#### 2.3.2. Protein estimation

The total protein concentration in the exosome samples was quantified using the Bradford assay (53), a colorimetric method based on the binding of Coomassie Brilliant Blue G-250 dye to proteins. Bovine serum albumin (BSA) was used as the reference standard for generating the calibration curve, with a concentration range of 20 to 150 μg /mL. The absorbance was measured at 595 nm, and protein concentrations in the samples were interpolated from the standard curve.

### 2.4. Characterization of exosomes

Exosomes purified through Octyl-sepharose and Sephadex-G-100 column chromatography were analyzed using transmission electron microscopy (TEM, Morgagni 268D at SAIF, AIIMS, New Delhi, India) to observe their characteristic cup-shaped morphology (54).

In addition, purified exosomes from serum and PF samples of RHD patients, CABG patients and healthy individuals were run on SDS (10%) gel (55). Proteins were transferred onto a polyvinylidene fluoride (PVDF) membrane. The blot was blocked with 5% (w/v) skimmed milk. Blot was incubated for 2 hours with biotin-labeled anti-human CD63 (Biolegend; 1:500) primary antibody. Following five washes, the blot was incubated at room temperature for two hours with Streptavidin-HRP-conjugated peroxidase (Biolegend; 1:10,000) secondary antibody, prepared with skim milk (2% w/v; 10 mL). After five washes with TBS-T buffer, protein bands were visualized using enhanced chemiluminescence (ECL; Bio-Rad) western blotting substrate. The blots were then examined by exposing them in a darkroom with the Protein simple fluorchem M imaging system (Bio-Techne).

### 2.5. Effect of purified exosomes on HUVECs

#### 2.5.1. HUVECs culture

HUVECs were isolated from umbilical cords obtained from healthy mothers at the Obstetrics and gynecology (OBGY) Department, PGIMER, Chandigarh, India (56, 57). Immediately following delivery, the umbilical cord was placed in transfer buffer (1X phosphate-buffered saline with penicillin; 50 mL) on ice. The outer surface of the cord was cleaned with ethyl alcohol (70% v/v), then rinsed with PBS (1X). The ends of the cord were trimmed using a sterile surgical blade, and a cannula attached to a syringe was inserted at one end. The umbilical vein was flushed with PBS *via* syringe to remove erythrocytes. After clamping the cord ends with a sterile hemostat, the umbilical vein was filled with a collagenase type II solution (0.2% w/v; Gibco, Thermo fisher scientific) and incubated at room temperature for 20 minutes. Thereafter, collagenase solution containing HUVECs was collected from the umbilical vein and centrifuged (1200 rpm x 5 minutes) to pellet the cells. HUVECs were cultured in endothelial cell growth medium (Cell applications, INC) supplemented with 10% (v/v) fetal bovine serum (FBS), and 1X antibiotic antimycotic solution (Himedia). The HUVECs, were maintained in a tissue culture flask (T25) under a humidified environment (95%) at 37^°^C with 5% CO_2_. When HUVECs obtained 80% confluence they were subsequently, trypsinized (0.25% w/v trypsin; 500 µL). Thereafter 0.5x 10^6^ cell/well were seeded in 6 well plate and subjected to treatments (100 μg), including serum/ PF, serum/ PF purified exosomes and combinations of serum with serum exosomes/ PF with PF exosomes obtained from RHD patients (HUVECs *in-vito* RHD model) and CABG patients. As a comparative measure, HUVECs were also treated with serum and purified serum exosomes from healthy individuals, and a control/placebo solution containing 10 mM phosphate buffer, pH 8.0. The cellular morphology was observed using an inverted-microscope (Olympus CKX41, Japan).

#### 2.5.2. cDNA synthesis

Transfected HUVECs were trypsinized, and the resulting cell pellets were collected by centrifugation at 1200 rpm for 10 minutes. Total RNA was extracted using 500 μL of TRIzol reagent. RNA concentration (ng/µL) and purity were determined by measuring the A260/A280 ratio, which typically ranged from 1.8 to 2.0 (58). cDNA was synthesized from 300 ng of total RNA using a high-capacity cDNA synthesis kit, (**Table 1**) and stored at −20°C until further use.

**Table 1:** Components used for cDNA synthesis

| Components | Volume (μL) |
| --- | --- |
| Nuclease free water | X μL |
| 10X RT Buffer | 2.0 μL |
| 20X dNTP mix (100 mM) | 0.8 μL |
| 10X RT Random primers | 2.0 μL |
| Reverse transcriptase | 1.0 μL |
| Total RNA | 300 ng |
| Total volume | 20 μL |

#### 2.5.3. qRT-PCR analysis

The quantitative reverse transcription polymerase chain reaction (qRT-PCR) was performed using Roche light-cycler® 96 and Biorad CFX96 systems to evaluate the relative expression levels of VEGF-A, RASA-1, IL4, endostatin, DLL-4, MMP2, and TP53. For the analysis, 1X SYBR green (Thermo Fisher Scientific) was used with gene-specific primers (**Table 2**). The fold change in mRNA levels between the test and control samples were calculated using the 2^-ΔΔCt^ formula (59). Reaction conditions for qRT-PCR gene(s) expression analysis are provided (**Table 3**).

**Table 2:**
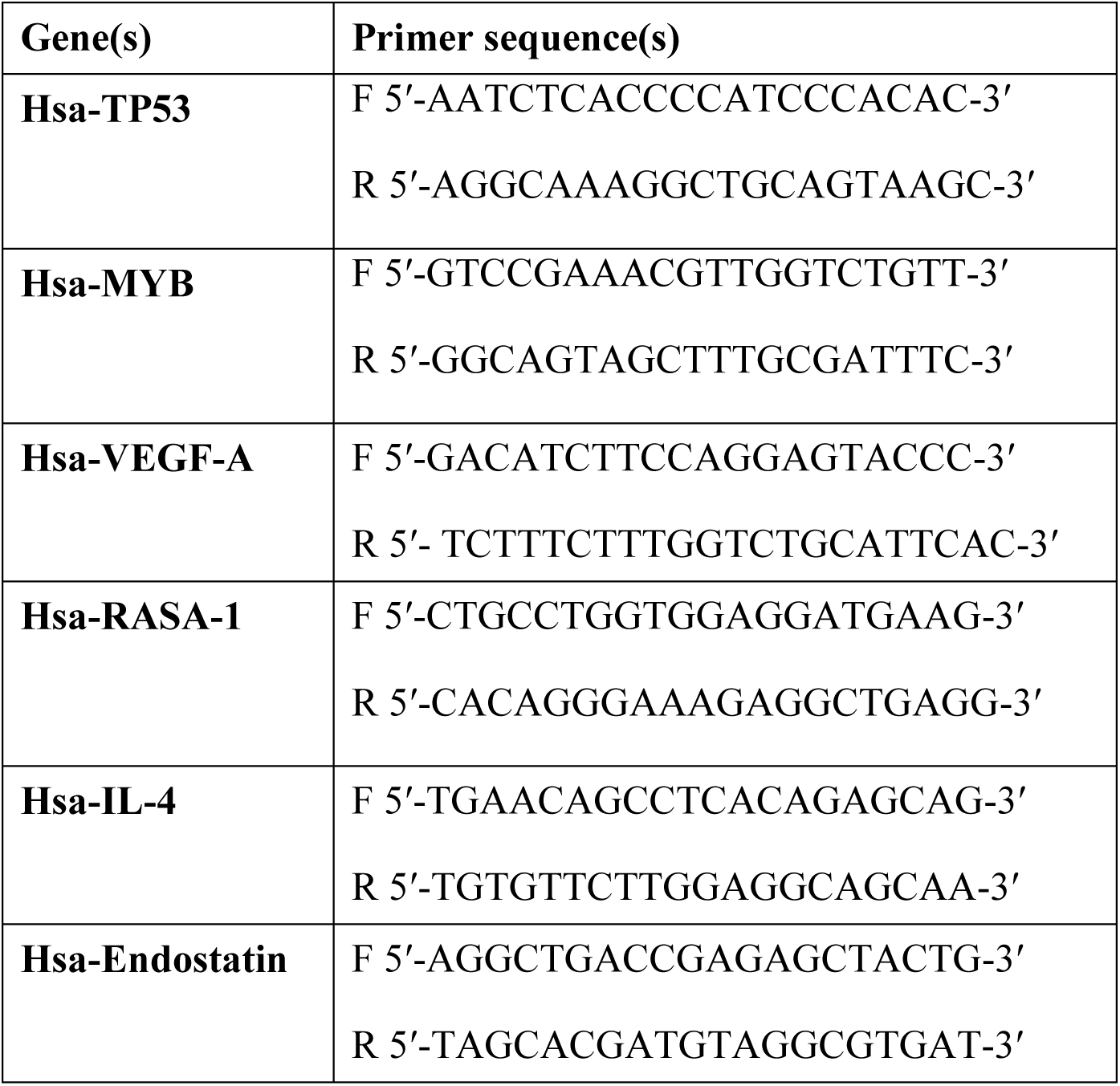
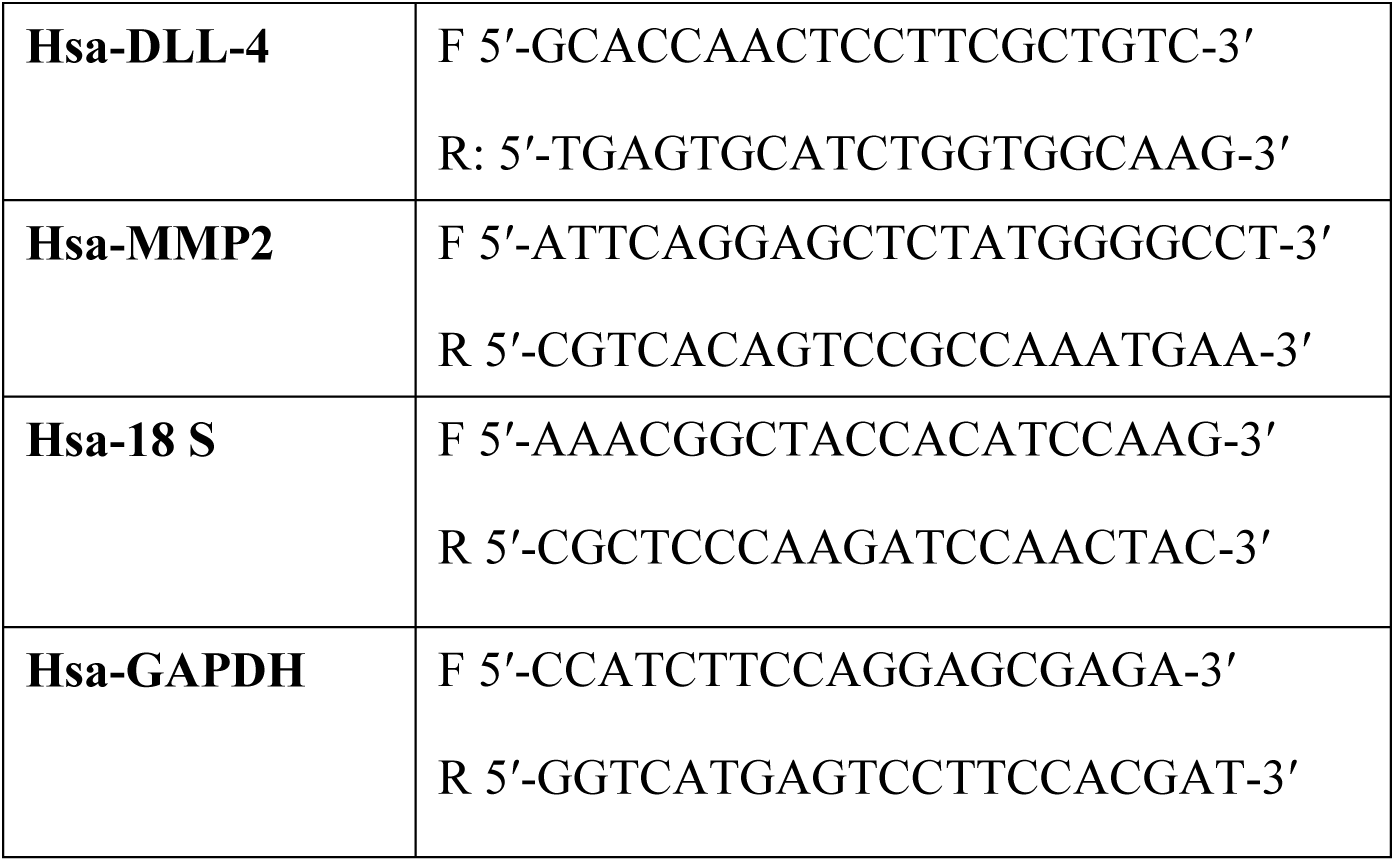
Oligonucleotide sequences of primers used for angiogenesis modulatory gene(s) expression

**Table 3:**
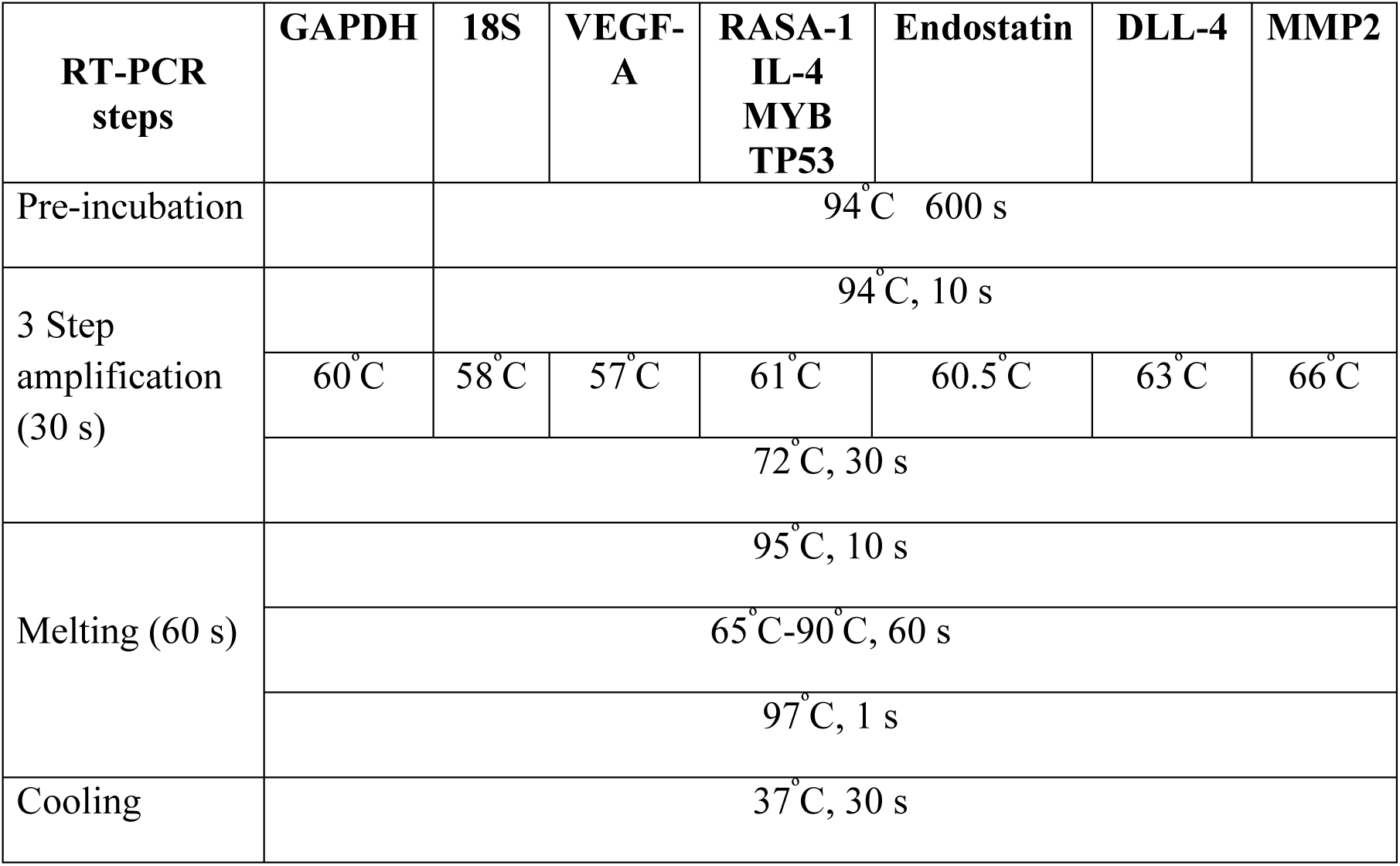
Reaction conditions used for angiogenesis modulatory gene(s) expression analysis

Where,

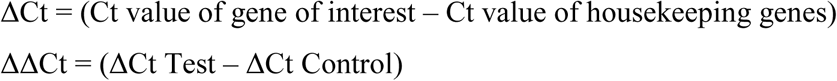

### 2.6. Expression of dysregulated miR-150-5p

A previous study from our lab showed, enhanced expression of miR-150-5p in serum exosomes from RHD patients, as compared to those from healthy individuals, based on NGS analysis (57). Total RNA including miRNAs were isolated from serum and PF exosome samples by Trizol method. RNA (ng/µL) concentration were assessed by measuring absorbance using a multi-mode reader (Infinite Pro 200 Tecan). Subsequently, miRNA underwent a two-step process: First, polyadenylation, followed by cDNA synthesis using the PCR system (Applied biosystems). For polyadenylation, the components were gently mixed in a microcentrifuge tube and incubated at 37^°^C for 30 minutes (**Table 4**). The polyadenylation reaction was terminated by incubating at 95^°^C for 5 minutes. For cDNA synthesis from the polyadenylated samples, the components (**Table 5**), were gently mixed and initially incubated at 55^°^C for 5 minutes, then incubated at 25^°^C for 15 minutes. Reverse transcription reaction was performed (42^°^C x 40 minutes), followed by 5 minutes incubation at 95^°^C to terminate the reaction and cDNA was stored at -20^°^C for future use. In column chromatography purified serum exosomes of RHD patient’s and healthy individuals (n= 20 each) miR-150-5p expression were evaluated. The miR- 150-5p-specific primers, along with the U6 snRNA primer, were used for the expression analysis (**Table 6**). qRT-PCR was performed on the light-cycler^®^ 96 (Roche, Germany). The reaction mixture consisted of 5 µL of 1X SYBR green (Thermo fisher scientific), forward and reverse primers (0.5 µL each), cDNA template (1.0 µL), and nuclease-free water (3.0 µL) and amplification was performed for 40 cycles (**Table 7**). The miR-150-5p relative expressions were calculated in samples by 2^−ΔCT^ formula.

**Table 4:** Components used for poly-adenylation of miRNA

| Components | Volume |
| --- | --- |
| Nuclease free water | X μL |
| 5X PolyA polymerase buffer | 4 μL |
| ATP | 1 μL |
| *E.Coli* polyA Polymerase | 1 μL |
| RNA template | 150 ng |
| Total volume | 20 μL |

**Table 5:**
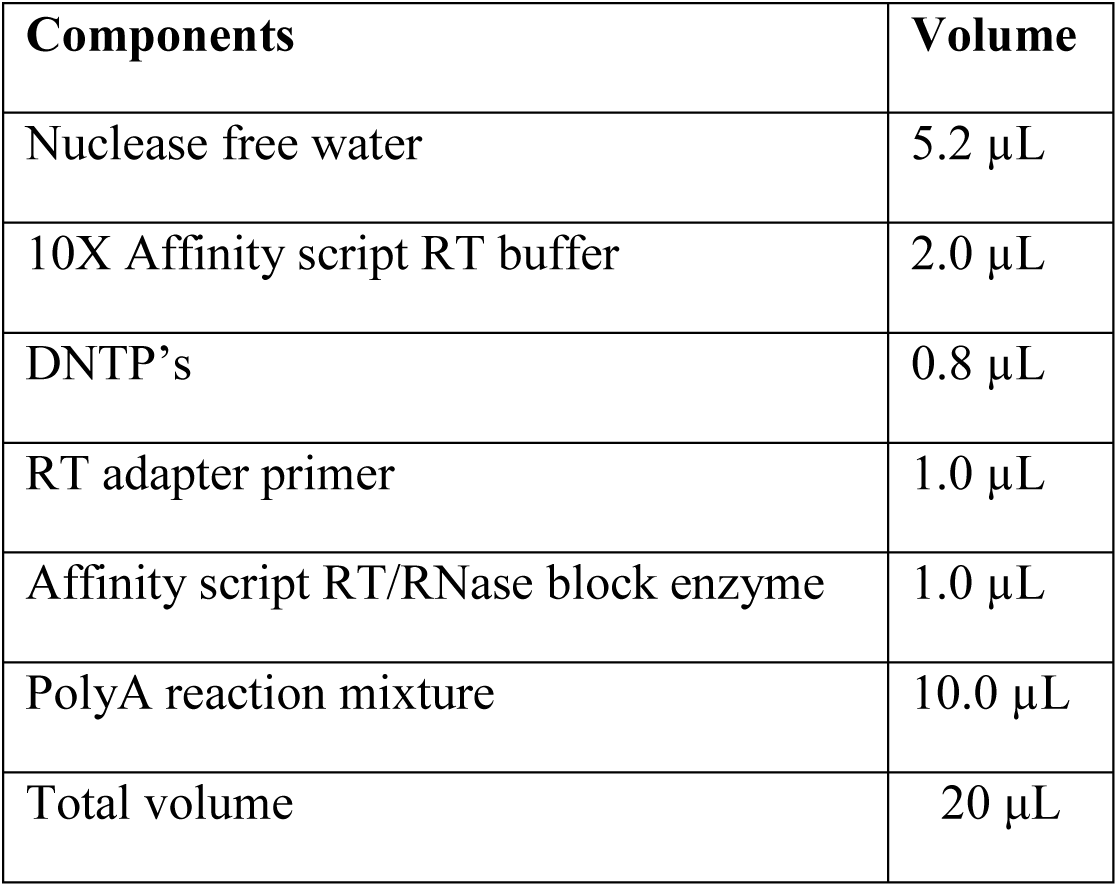
Components used for miRNA to cDNA synthesis

| Components | Volume |
| --- | --- |
| Nuclease free water | 5.2 μL |
| 10X Affinity script RT buffer | 2.0 μL |
| DNTP's | 0.8 μL |
| RT adapter primer | 1.0 μL |
| Affinity script RT/RNase block enzyme | 1.0 μL |
| PolyA reaction mixture | 10.0 μL |
| Total volume | 20 μL |

**Table 6:**
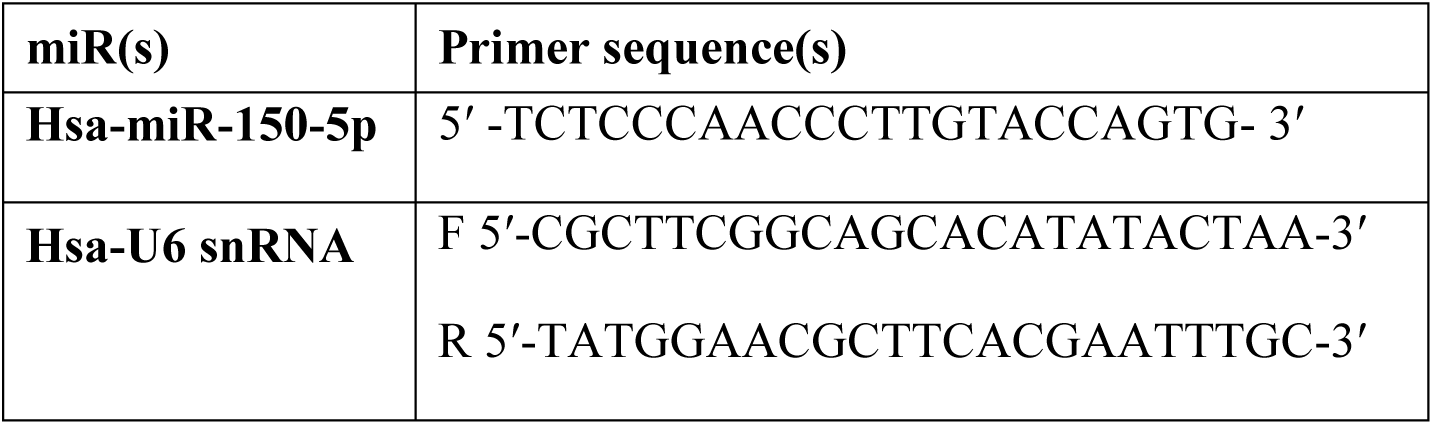
Oligonucleotide sequences of primers used for miRNA expression

**Table 7:** Reaction conditions used for of hsa-miRs and U6 for real time PCR

| **RT-PCR steps** | **U6**<br>**snRNA** | **miR-150-5p** |
| --- | --- | --- |
| Pre incubation | 94°C, 600 s |  |
| Amplification | 94°C, 10 s |  |
|  | 56.0°C, 20 s | 61.0°C, 20 s |
|  | 72°C, 20 s |  |
| Melting | 95°C, 10 s |  |
|  | 61°C, 60 s |  |
|  | 97°C, 1 s |  |
| Cooling | 37°C, 30 s |  |

Where,

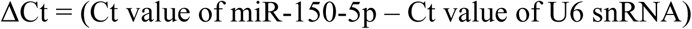

*In-silico* analysis was performed to identify the target genes of miR-150-5p that regulate angiogenesis pathways. A review of literature and *in-silico* predictions revealed MYB and TP53 are potential target genes of miR-150-5p, which may modulate angiogenesis. Therefore, gene-specific primers for MYB and TP53 were designed to assess their relative expression as selected targets of miR-150-5p.

### 2.7. Angiogenic properties of exosomes-induced HUVECs

#### 2.7.1. Effect of miR-150-5p mimics/ anti-miR-150-5p

Hsa-miR-150-5p mimics and hsa-anti-miR-150-5p were purchased from Thermo fisher scientific and used for transfection. HUVECs (2 x 10^5^ cells/ well) were seeded in 24-well plate with antibiotic free low serum media (5% FBS). Cultured HUVECs were treated with serum exosomes from RHD and healthy individual’s serum exosomes (20 µg each), along with transfections of miR-150-5p mimics (30 nM), anti-miR-150-5p (30 nM), and suramin (40 µM) using lipofectamine (Thermo Fischer Scientific). After treatment, HUVECs were observed for morphological changes, and the images were analysed using an inverted microscope.

#### 2.7.2. Angiogenesis assay

HUVECs (2 x 10^4^ cells per well) were plated onto an extracellular matrix gel (Angiogenesis assay kit; Abcam) in endothelial cell growth medium (containing 5% v/v FBS without antibiotics) in a 96-well plate. The cultured cell were then treated with purified exosomes, transfected with miR-150-5p mimics (30 nM), anti-miR-150-5p (30 nM), and suramin (40 µM) using lipofectamine. After transfection, the cells were cultured for 15 hours at 37^°^C with 5% carbon dioxide, followed by the removal of the medium and staining with staining dye (1:200; 100 µL per well). The formations of endothelial angiogenic tube-like structures were examined using fluorescence microscopy.

### 2.8. Statistical analysis

Statistical analysis was performed with GraphPad prism v.8. ANOVA was applied to evaluate group differences, followed by Dunnett’s multiple comparisons test to compare each group to the control HUVECs group. An unpaired t-test was used to assess relative miR-150-5p and TP53 expression. Data are presented as mean ± SEM.

## 3. RESULTS

### 3.1. Exosome isolation and characterization from RHD patient samples: Selection of column chromatography technique

Blood and PF samples were collected from confirmed RHD patients (10 females, 10 males), as well as PF samples from age and sex matched CABG patients (10 females, 10 males). Additionally, blood samples were obtained from age and sex-matched healthy individuals (10 females, 10 males). Pooled serum sample from RHD patients was subjected to fractionation using DEAE-cellulose and Octyl-sepharose columns. Twenty fractions (2 mL each) were collected, and their absorbance was measured at 450 nm. The elution profile revealed distinct peaks for biomolecules in the DEAE-cellulose column (fractions 2-4; **Fig. 1.A**) compared to the Octyl-sepharose column (fractions 5-6; **Fig. 1.B**). Among the eluted fractions, those with the highest absorbance peaks at 450 nm (*i.e.,* the 3^rd^ fraction from the DEAE-cellulose column, 5^th^ and 13^th^ fractions from the Octyl-sepharose column) were selected for TEM analysis. TEM results showed a limited presence of exosomes in the selected fractions from the DEAE-cellulose column (3^rd^ fraction, **Fig. 1.C**) and Octyl-sepharose column (5^th^ fraction, **Fig. 1.D**). In contrast, abundant exosomes were observed in the selected fraction isolated by Octyl-sepharose column chromatography (13^th^ fraction, **Fig. 1.E**). Based on these findings, Octyl-sepharose column chromatography was selected as the preferred method for exosome isolation.

**Figure 1:**
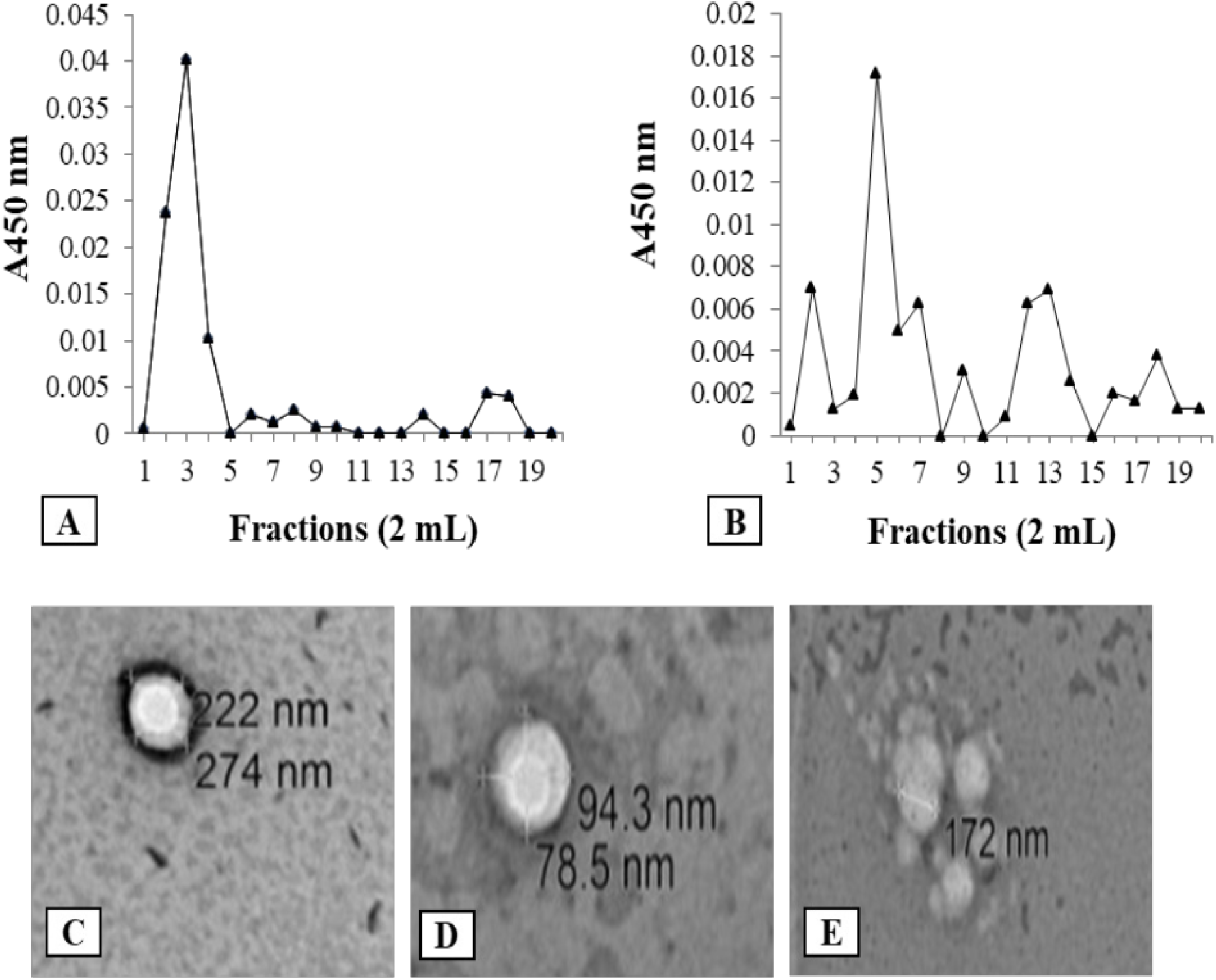
Elution profiles and transmission electron microscopy (TEM) of purified exosomes from RHD patient samples: (A) Elution profile of biomolecules from Diethylaminoethyl-cellulose (DEAE-cellulose) column chromatography, showing fractionation with potassium phosphate buffer (50 mM, pH 8.0) with a NaCl gradient (0.5 M in fractions 1 to 10 and 1.0 M in fractions 11 to 20). (B) Elution profile from Octyl-sepharose column chromatography using potassium phosphate buffer (50 mM, pH 8.0) with an ammonium sulfate gradient (0.5 M in fractions 1-10, transitioning to no ammonium sulfate in fractions 11-20). (C) transmission electron microscopy (TEM) image of exosomes isolated from the 3^rd^ fraction of the DEAE-cellulose column (0.5 M NaCl), showing a limited presence of exosome-like vesicles. (D) TEM image of exosomes isolated from the 5^th^ fraction of the Octyl-sepharose column (potassium phosphate buffer, 0.5 M ammonium sulfate), showing limited presence of exosomes. (E) TEM image of exosomes isolated from the 13^th^ fraction of the Octyl-sepharose column (potassium phosphate buffer, no ammonium sulfate), demonstrating a high concentration of exosomes. Exosomes were stained with uranyl acetate and visualized under TEM. These results indicate that Octyl-sepharose column chromatography provides a more efficient method for exosome isolation and purification from biological fluids.

### 3.2. Isolation and characterization of exosomes

The pooled PF and serum samples from RHD patients were subjected to fractionation using Octyl-sepharose column chromatography. Among the eluted fractions, the 3^rd^, 4^th^, and 9- 12^th^ fractions, which showed the highest absorbance at 280 nm (**Fig. 2.A**), were pooled, desalted, and concentrated using ultracentrifugal filters (10 kDa). The resulting concentrated samples underwent further fractionation using Sephadex G-100 column chromatography. Fractions exhibiting Ach-activity, particularly those from the fifth to ninth fractions of Sephadex G-100 column chromatography (**Fig. 2.B**), were stored at -20^°^C. The Ach-colorimetric assay revealed a yellow coloration (**Fig. 2.C**). The western blot analysis showed a 30 kDa protein band corresponding to CD-63, exosome marker (**Fig. 2.D**). TEM analysis revealed the distinct cup-shaped morphology in the column chromatography purified exosome-containing fractions (**Fig. 2.E-I**). The combined results from TEM and the CD-63 western blotting confirmed the observed structures as exosomes.

**Figure 2:**
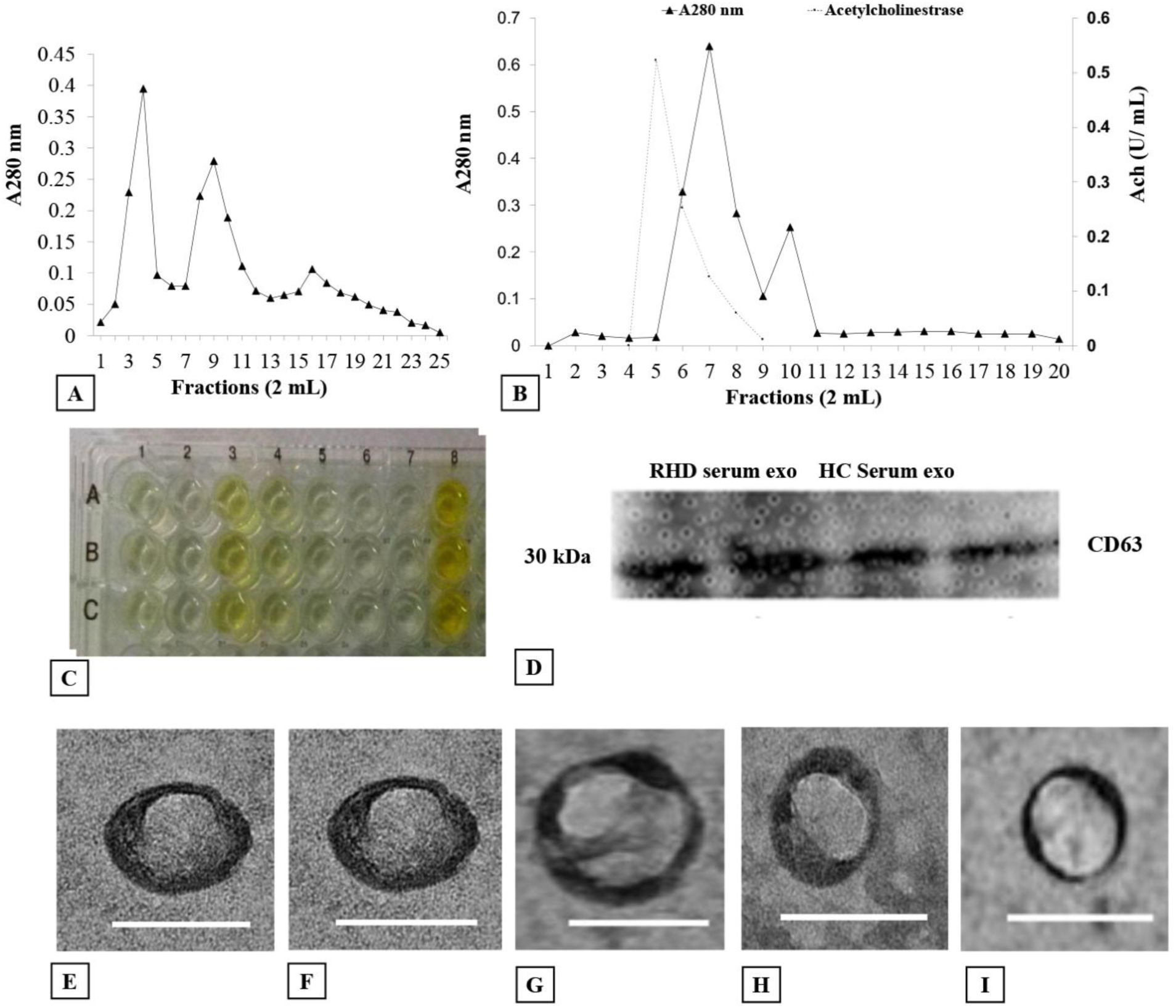
Elution profile, colorimetric assay, and characterization of column-purified exosomes: (A) Elution profile of exosomes obtained through Octyl-sepharose column chromatography using potassium phosphate buffer (50 mM, pH 8.0). The absorbance at 280 nm was measured for each fraction to identify peaks corresponding to biomolecules. (B) Sephadex G-100 column chromatography profile showing the absorbance at 280 nm (solid line) and acetylcholinesterase (Ach) activity (dashed line) in the eluted fractions. (C) Acetylcholinesterase colorimetric assay on a 96-well plate to identify Ach activity in exosome-containing fractions. Lane 1 represents the blank, Lanes 2-7 correspond to Sephadex G-100 purified fractions (5^th^ to 10^th^), and Lane 8 represents the Octyl-sepharose-purified and concentrated exosomes. (D) Western blot analysis of exosomes for the CD63 exosome surface marker, using samples from RHD patient exosomes purified via column chromatography. The CD63 band confirms the identity of the structures as exosomes. (E-I) Transmission electron microscopy (TEM) images (scale bar = 200 nm) of exosomes stained with phospho-tungstic acid, isolated from various sources: Serum/PF from RHD patients, Serum/PF from CABG patients and healthy control serum. TEM analysis reveals characteristic cup-shaped morphology of the exosomes. All results combined demonstrate successful isolation, purification, and characterization of exosomes.

### 3.3. Effect of RHD patient-derived exosomes on HUVECs

HUVECs were isolated from umbilical cords obtained from healthy mothers at the Obstetrics and gynecology (OBGY) Department, PGIMER, Chandigarh, India. HUVECs were cultured and maintained in endothelial cell growth medium enriched with 10% v/v FBS. Upon reaching 50-60% confluence, the HUVECs were treated with (100 μg) serum or PF, serum exosomes and combinations of serum with serum exosomes or PF exosomes with PF from RHD patients and CABG patients. For comparison, HUVECs were also treated with serum and purified serum exosomes obtained from healthy individuals. After 24 hours, the morphological features of the treated HUVECs (at 70-80% confluence, passage no. 4) and the control group were examined (Supplementary, **Fig. S.1**). RNA was extracted from transfected HUVECs using the trizol method, followed by the cDNA synthesis, to assess the gene expression of VEGF-A (angiogenic), IL-4 (inflammatory), RASA-1 (angiogenesis inhibitor), endostatin (anti-angiogenic), DLL-4 (anti-angiogenic), MMP2 (angiogenic-switch) and TP53 (miR-150-5p target gene).

#### 3.3.1. Gene expression of VEGF-A, IL-4, and RASA-1 in HUVECs treated with PF and serum exosomes from RHD patients, CABG patients or healthy controls

To investigate the angiogenic potential of circulating exosomes from RHD patients, CABG patients or healthy control individual’s serum and PF we assessed the expression of angiogenesis key genes in HUVECs following treatment with exosomes. A significant up- regulation of VEGF-A (2.73-fold increase; p= 0.0002), was observed in HUVECs treated with CABG PF-derived exosomes, followed by HUVECs treated with RHD serum exosomes (2.70- fold, p= 0.0002) and RHD serum (2.5-fold; p= 0.001), suggesting angiogenic stimulus (**Fig. 3.A**). In contrast, the pro-inflammatory cytokine IL-4 (15.17-fold; p= 0.0001), exhibited the highest expression in HUVECs treated with RHD serum-derived exosomes, indicating a strong inflammatory response (**Fig. 3.B**). Interestingly, the expression of RASA-1, a known angiogenesis inhibitor, was markedly upregulated in HUVECs treated with CABG-PF (14.85- fold; p= 0.0001), followed by CABG serum-derived exosomes (13.25-fold; p= 0.0001), indicating a potential anti-angiogenic regulation in this group. Conversely, RASA-1 expression was downregulated (0.65-fold; p= 0.999) in HUVECs treated with RHD serum exosomes (**Fig. 3.C**). Overall, these results highlight that the upregulated VEGF-A (angiogenic) expression and IL-4 (inflammatory) gene expression, whereas reduced RASA-1 (angiogenesis inhibitor) gene expression profiles induced by exosomes from RHD patients, suggesting that the exosomes cargo significantly influence their regulatory effects on endothelial cells.

**Figure 3:**
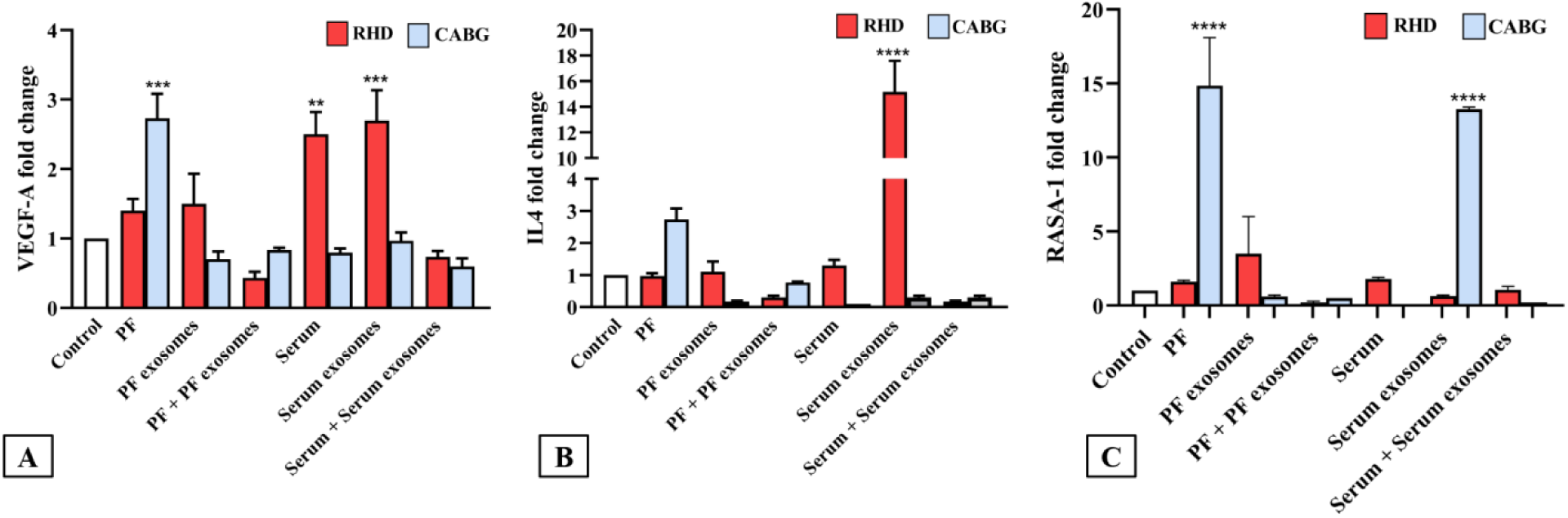
Gene Expression of angiogenesis modulating genes in HUVECs treated with exosomes from RHD and CABG patients: (A) Relative gene expression of vascular endothelial growth factor-A (VEGF-A), a key angiogenic gene, in human umbilical vein endothelial cells (HUVECs) treated with serum or PF exosomes from RHD and CABG patients. (B) Relative gene expression of interleukin 4 (IL-4), an inflammatory cytokine, in HUVECs treated with exosomes. (C) Relative gene expression of RAS p21 protein activator 1 (RASA-1), an angiogenesis inhibitor, in the same HUVEC treatment groups. The data were analyzed using Dunnett’s multiple comparisons ANOVA, and statistical significance is indicated by * where * P < 0.05, ** P < 0.01, *** P < 0.001, and **** P < 0.0001 compared to control HUVECs. All data are presented as mean ± SEM. Serum exosomes from RHD patients significantly upregulate VEGF-A and IL-4 expression, while downregulating RASA-1, indicating a potential angiogenic and inflammatory effect of RHD patients serum-derived exosomes on HUVECs.

#### 3.3.2. Comparative gene expression of Endostatin, DLL-4 and MMP2 in HUVECs treated with RHD patients, CABG patients or healthy control derived serum exosomes

To further investigate the effect of serum-derived exosomes in angiogenesis, we analyzed the expression of endostatin, DLL-4, and MMP2 in HUVECs treated with serum exosomes from RHD patients, CABG patients and healthy controls. Endostatin, an anti-angiogenic factor, was significantly upregulated (3.73-fold; p= 0.0001) in HUVECs treated with serum exosomes from healthy individuals compared to control cells. In contrast, HUVECs treated with RHD serum exosomes showed a reduced endostatin expression (0.4-fold; p= 0.070; **Fig. 4.A**). DLL-4 (anti-angiogenic), showed the highest expression (2.13-fold; p= 0.0015) in HUVECs treated with healthy serum exosomes, followed by CABG serum exosomes (1.63-fold; p= 0.0382) and RHD serum exosome treated group (1.4-fold; p= 0.0201; **Fig. 4.B**). In contrast, expression of MMP2, a marker associated with the angiogenic switch, was significantly elevated in HUVECs exposed to RHD serum exosomes (1.53-fold; p= 0.0312) relative to control cells (**Fig. 4.C**). MMP2 expression was not significantly altered in the healthy or CABG exosome-treated groups. These results suggest that serum exosomes from healthy individuals favor anti-angiogenic signaling, as indicated by increased endostatin and DLL-4 expression, whereas RHD-derived exosomes promote a pro-angiogenic environment, reflected by reduced anti-angiogenic endostatin gene expression and upregulation of MMP2.

**Figure 4:**
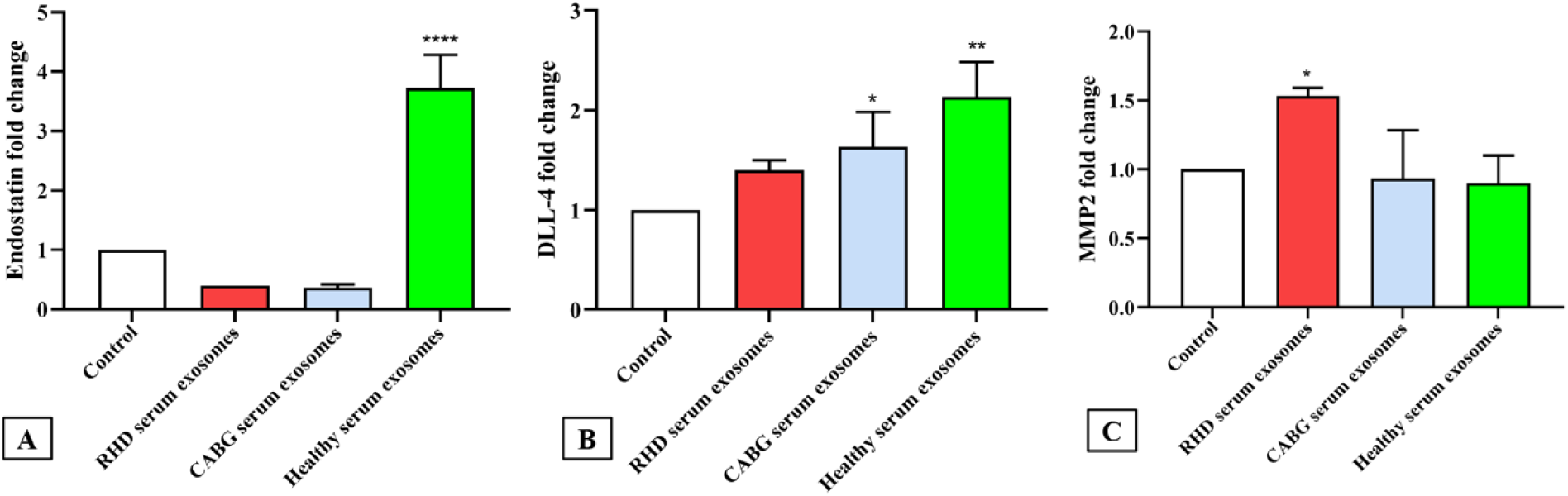
Expression levels of genes associated with angiogenesis in HUVECs treated with serum exosomes from RHD, CABG, and healthy individuals: (A) Relative gene expression of endostatin, an anti-angiogenic factor, in human umbilical vein endothelial cells (HUVECs) treated with serum exosomes from RHD, CABG patients, and healthy individuals. (B) Relative expression of Delta-like 4 (DLL-4), an anti-angiogenic gene, in the same treatment groups. (C) Relative expression of matrix metalloproteinase-2 (MMP2), a gene associated with angiogenic switch, in HUVECs treated with exosomes. Data were analyzed using Dunnett’s multiple comparisons ANOVA, with statistical significance indicated by * where * P < 0.05, ** P < 0.01, *** P < 0.001, and **** P < 0.0001 compared to control HUVECs. All data are presented as mean ± SEM. These findings show that serum exosomes from healthy individuals significantly upregulated endostatin and DLL-4, while serum exosomes from RHD patients downregulated endostatin and upregulated DLL-4 and MMP2, suggesting a shift toward angiogenesis.

### 3.4. miR-150-5p expression in serum exosomes from RHD patients: Targeting TP53

The purified pooled serum exosomes from RHD patients exhibited a more pronounced angiogenic effect on HUVECs compared to PF exosomes. A previous study from our lab found, enhanced expression of miR-150-5p in serum exosomes of RHD patients, as identified by NGS, compared to healthy individuals (57). Based on these findings, miR-150-5p was selected for further study. We observed a significant upregulation of miR-150-5p expression in serum exosomes (34.29 ± 9.524-fold mean difference; p= 0.0007; **Fig. 5.A**) from RHD patients compared to those from healthy individuals serum exosomes, with U6 snRNA as an internal control. miRDB predicted Myb-related protein B (MYB) and tumor protein (TP53) as potential targets of miR-150-5p (Supplementary, **Fig. S.2**). Furthermore, significant downregulation of TP53 (0.227-fold; p= 0.006; **Fig. 5.B**) gene expression was observed in HUVECs treated with serum exosomes from RHD patients.

**Figure 5:**
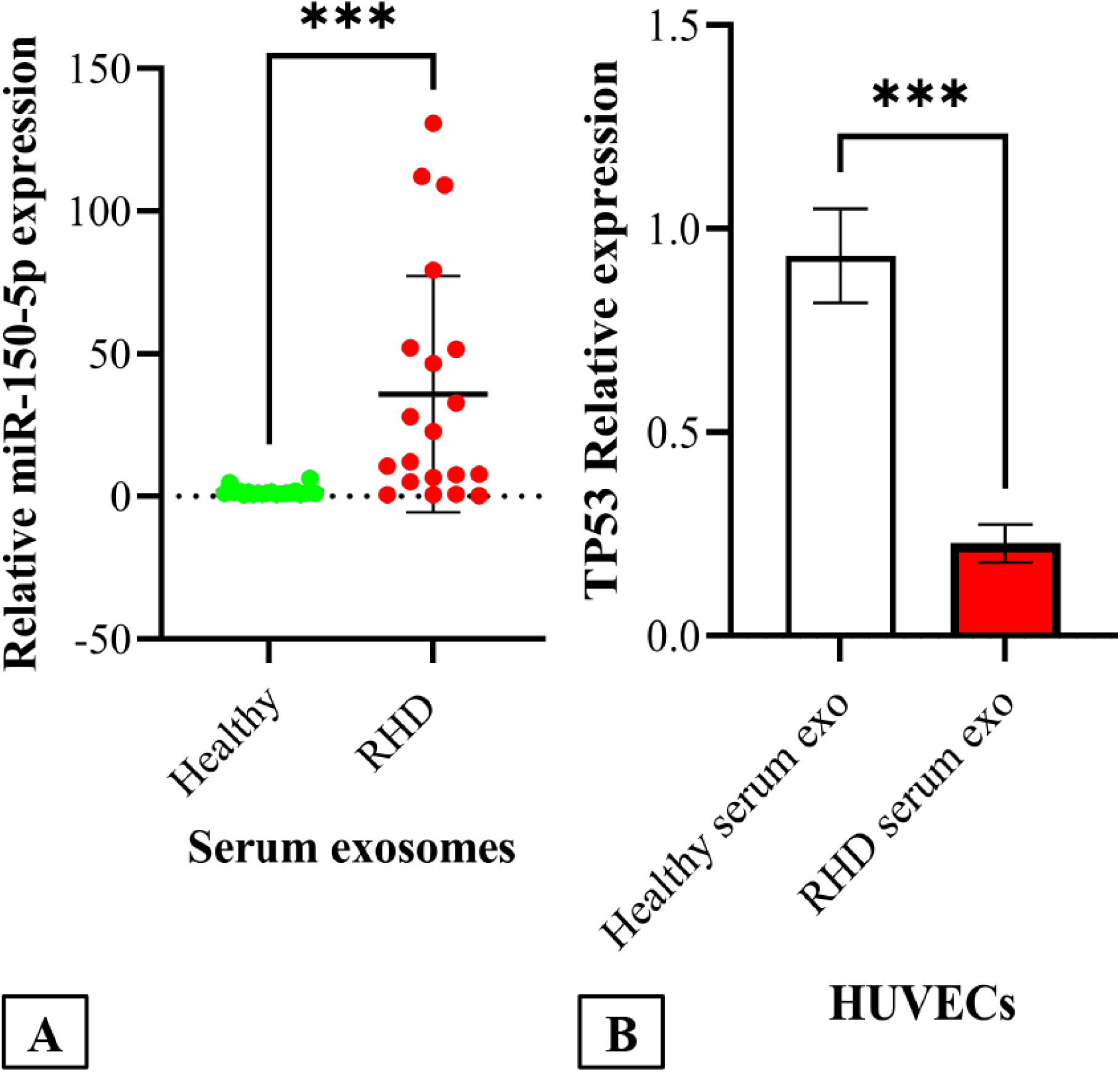
Expression of miR-150-5p in serum exosomes from RHD patients and TP53 in HUVECs treated with exosomes: (A) Relative gene expression levels of miR-150-5p in serum exosomes from RHD patients and healthy individuals, with U6 snRNA used as an internal control. The data show a significant upregulation of miR-150-5p in exosomes from RHD patients (33.80 ± 13.2-fold, P = 0.0162) compared to healthy controls. (B) Gene expression of TP53 in HUVECs treated with serum exosomes from RHD patients, revealing significant downregulation (0.267-fold, P = 0.006) of TP53, a predicted target of miR-150-5p. Statistical analysis was performed using Dunnett’s multiple comparisons ANOVA, and data are presented as mean ± SEM. Significant P-values are indicated by * where *** P < 0.001 compared to control. These findings suggest that exosomes from RHD patients may influence endothelial cell function through miR-150-5p-mediated regulation of TP53.

### 3.5. Regulation of angiogenesis in HUVECs by RHD serum exosomes and miR-150-5p modulation

HUVECs were isolated from umbilical cord and maintained in the laboratory. The cells were then treated with purified exosomes and transfected with miR-150-5p mimics (for miR- 150-5p overexpression) and anti-miR-150-5p (for miR-150-5p suppression) using Lipofectamine. As compared to the control HUVECs (**Fig. 6.A**), stimulation with pooled serum exosomes (20 µg) from RHD patients (**Fig. 6.B**), CABG patients (**Fig. 6.C**) resulted in enhanced network formation, and the development of cage-like structures. Serum exosomes from both RHD and CABG patients demonstrated enhanced angiogenic potentials in HUVECs, in contrast to serum exosomes from healthy individuals (**Fig. 6.D**). Additionally, HUVECs transfected with 30 nM miR-150-5p mimics (**Fig. 6.E**) showed enhanced angiogenic potential, compared to cells transfected with anti-miR-150-5p (**Fig. 6.F**) and treated with 40 µM suramin (**Fig. 6.G**).

**Figure 6:**
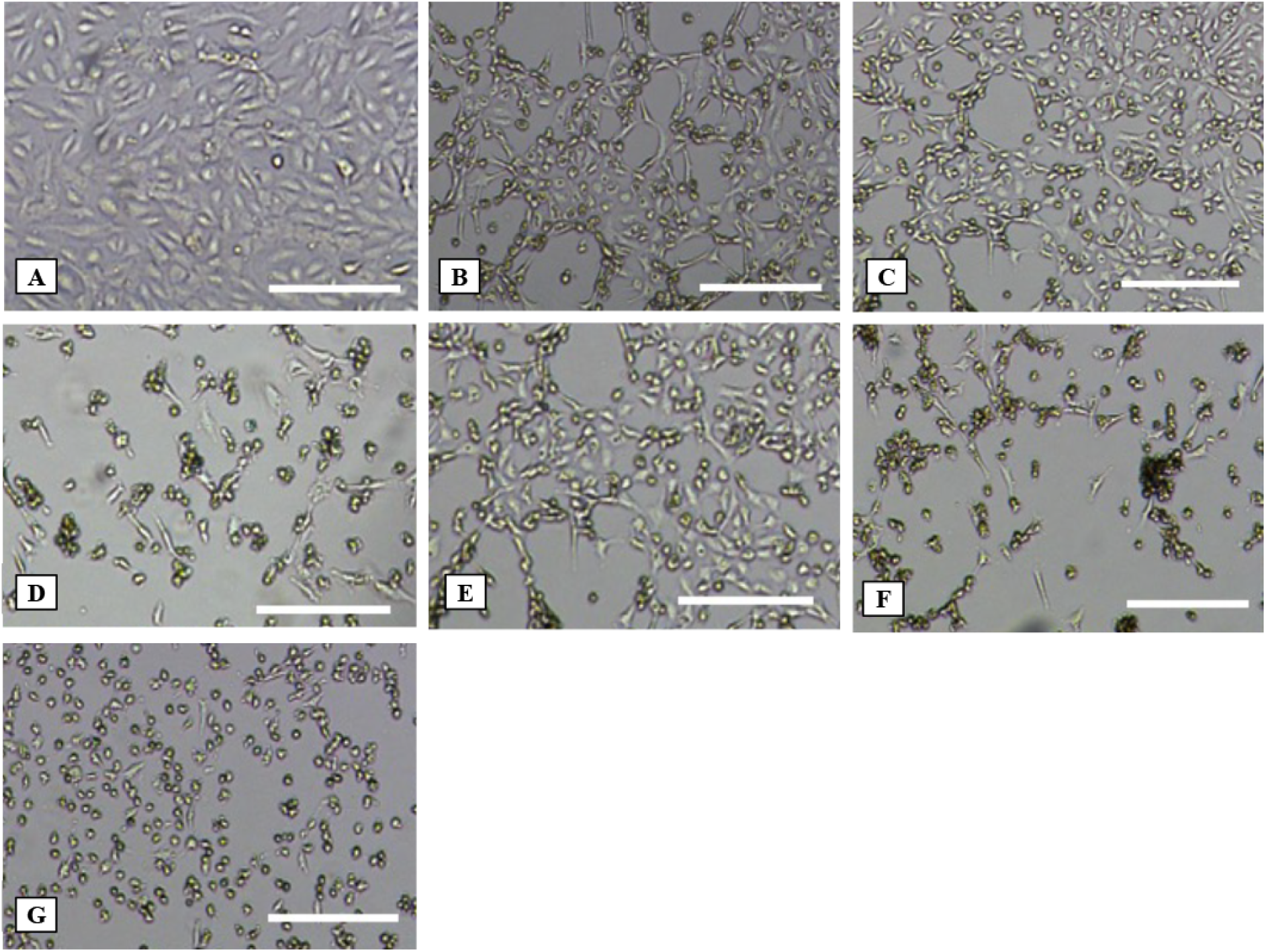
Angiogenic effects of hsa-miR-150-5p mimics and serum exosomes on HUVECs: (A-H) Morphological changes in HUVECs treated with different conditions, observed under an inverted microscope at 40X (4X x 10X= 40X magnifications, scale bar = 200 µm) magnification. (A) Control HUVECs show normal growth patterns. (B) Serum exosomes from RHD patients (20 µg) promote enhanced network formation with cage-like structures. (C) Serum exosomes from CABG patients (20 µg) also stimulate network formation with cage-like structures. (D) Serum exosomes from healthy individuals (20 µg) does not show angiogenic effects and lead to cell death. (E) HUVECs transfected with 30 nM hsa-miR-150-5p mimics exhibit enhanced angiogenesis, with improved network formation. (F) Cells transfected with 30 nM hsa-anti-miR- 150-5p show reduced angiogenic potential and induced cell death. (G) HUVECs treated with 40 µM suramin lead to cell death. These results demonstrate that serum exosomes from RHD and CABG patients, as well as hsa-miR-150-5p mimics, enhance angiogenesis in HUVECs, while anti-miR-150-5p or suramin inhibit this effect.

### 3.6. Endothelial tube formation induced by RHD serum exosomes and miR-150-5p modulation

Angiogenesis assay kit was used to evaluate endothelial tube formation, which was observed through fluorescence microscopy. As compared to control HUVECs (**Fig. 7.A**), HUVECs treated with 20 µg of serum exosomes from RHD patient (**Fig. 7.B**), CABG patients (**Fig. 7.C**), showed elongation and the formation of angiogenic tube-like structures. In contrast, serum exosomes from healthy individuals did not promote tube formation and instead induced cell death (**Fig. 7.D**). Furthermore, transfection with 30 nM hsa-miR-150-5p mimics enhanced networking, and angiogenic tube formation in HUVECs (**Fig. 7.E**), as compared to those transfected with 30 nM hsa-anti-miR-150-5p (**Fig. 7.F**) or 40 µM suramin (**Fig. 7G**) which lead to cell death

**Figure 7:**
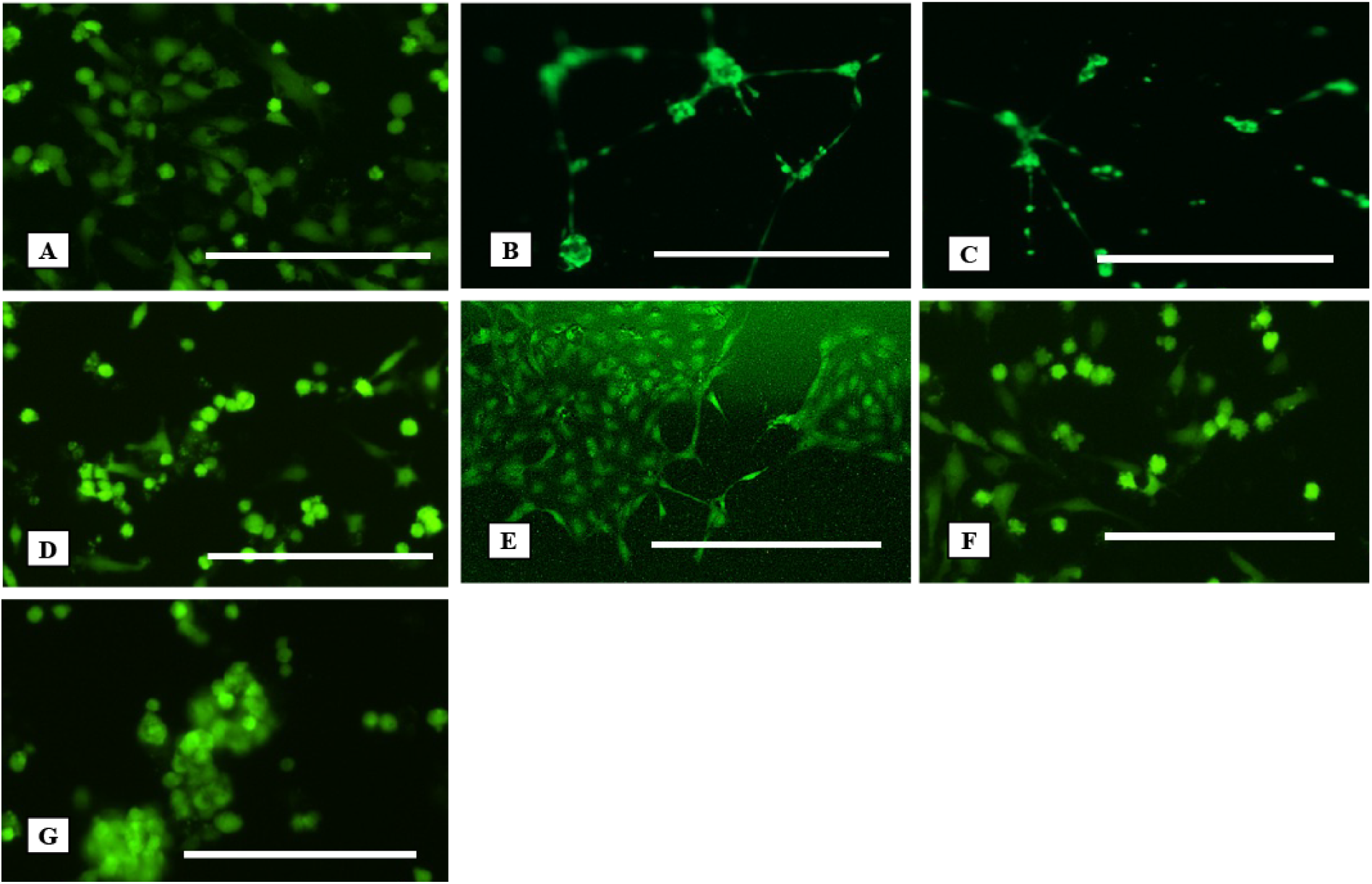
Induction of endothelial tube formation in HUVECs by serum exosomes from RHD and CABG patients. Role of miR-150-5p mimics: Endothelial tube formation assay was performed on HUVECs, observed via fluorescence microscopy at 40X (4X x 10X = 40 X magnifications, scale bar = 200 μm) magnification. (A) Control HUVECs show standard morphology without enhanced tube formation. (B) HUVECs treated with serum exosomes from RHD patients (20 μg) exhibit elongated cells and clear angiogenic tube-like structures. (C) HUVECs treated with serum exosomes from CABG patients (20 μg) show similar tube formation and network development. (D) Serum exosome from healthy individuals (20 μg) does not induce tube formation and lead to cell death. (E) HUVECs treated with 30 nM hsa-miR-150-5p mimics exhibit network formation and tube-like structures. (F) HUVECs transfected with 30 nM hsa-anti-miR-150-5p show inhibited tube formation and induced cell death. (G) Treatment with 40 μM suramin completely suppresses endothelial tube formation and lead to cell death. These results highlight the role of miR-150-5p mimics and exosomes in promoting angiogenesis in HUVECs.

## 4. DISCUSSION

RHD, a serious autoimmune consequence of rheumatic fever, primarily damages the mitral valves (2). Globally, an estimated 55 million individuals are affected by RHD, contributing to nearly 360,000 deaths annually. The World Health Organization (WHO) emphasizes the urgent need to explore the pathogenic mechanisms underlying RHD to improve our understanding of its pathogenesis. In this study, we hypothesized that RHD, which leads to mitral valve damage, is associated with aberrant angiogenesis, and exosome play a key role in mediating this process. More specifically, we propose that dysregulated miRNA enclosed within exosomes may serve as potential biomarkers reflecting RHD-specific pathological mechanism. To explore this, we analyzed both PF and serum samples from RHD patients. PF was selected due to its close proximity to the heart and potential to reflect localized pathological alterations.

For comparison, PF samples from CABG patients were collected during thoracic surgery for grafting because they had no history of autoimmune disease were included as surgical controls. Additionally, serum samples from healthy individuals were included as control to represent baseline systemic expression levels. Our investigation primarily focused on exploring the role of exosomes in regulating angiogenesis during RHD pathogenesis. Exosomes were isolated, purified, characterized from the PF and serum of RHD and CABG patients, as well as healthy individuals. We used hydrophobic interaction and size-exclusion column chromatography for exosome isolation: a method that does not require ultracentrifuge and helps to preserve the structural integrity of the exosomes. Prior studies suggest that size-exclusion chromatography yields exosomes with greater functional stability compared to ultracentrifugation methods (60, 61). Exosomes bind to the hydrophobic matrix of the Octyl-sepharose column matrix (62). After pooling and concentrating the fractions, the samples were further purified using a Sephadex G- 100 column. Fractions containing exosome were identified using an acetylcholinesterase (AChE) colorimetric assay, which detects AChE activity a common marker of extracellular vesicles, including exosomes (63). Acetylthiocholine is hydrolyzed by acetylcholinesterase to produce thiocholine (64), which form a yellow-colored 5-thio-2-nitro-benzoicacid (65, 66) compound, when reacts with DTNB reagent. Positive fractions, indicated by a yellow color, were then verified by TEM, which confirmed the cup-shaped morphology of exosomes and by immunoblotting for CD-63. Collectively, these analyses confirmed the successful isolation of exosomes.

Next, we assessed the angiogenic potential of isolated exosomes. Angiogenesis, often observed during the progression of RHD, is driven by key factors such VEGF-A, integrins, chemokines, angiopoietins, and oxygen-sensing agents among others (15). These factors activate signaling pathway that lead to endothelial cell migration, extracellular matrix remodeling, and new blood vessel formation (67). Endothelial cells, which line the inner surface of blood and lymphatic vessels, regulate vascular permeability and immune cell trafficking (68). Endothelial cells also regulate blood pressure by controlling vasoconstriction and vasodilation. The entire cardiovascular system, including the heart valves and chambers, is lined by a monolayer of endothelial cells. Given their crucial role in cardiovascular homeostasis, we used HUVECs to model the angiogenic environment of RHD *in-vitro*. HUVECs, were isolated, cultured, and treated with serum or PF samples including exosomes from RHD, CABG patients and healthy individuals. HUVECs treated with serum-derived exosomes from RHD patients exhibited elongation and stretching, suggestive of angiogenic activation (69). To confirm this, we evaluated the expression of angiogenesis-related genes: VEGF-A, RASA-1, IL4, endostatin, DLL-4, and MMP2. VEGF-A, a master regulator of angiogenesis (13), activates downstream signaling pathways such as Ras-Raf-Erk and PI3K pathways, promoting endothelial proliferation (70). RASA-1 negatively regulates this process by inactivating RAS and suppressing Raf/MEK/ERK signaling cascade (70, 71, 72, 73). IL4 has been shown to enhance angiogenesis (74, 75), while endostatin inhibits VEGF-A-mediated endothelial cell migration (76–78). MMPs contribute to angiogenesis by degrading extracellular matrix (ECM) and vascular basement membrane, thus facilitating neovascularization, supporting endothelial cell migration and proliferation. Studies have shown that DLL4 plays a crucial role in the tip (which are motile and respond to various growth factors, including VEGF-A) or stalk cell (which maintain blood vessel integrity) (79, 80) conversion mediated by the Notch pathway. DLL4 on the tip cell binds to Notch-1 on the stalk cell, leading to its cleavage and release of the Notch-1 intracellular domain (NCID) (81). NCID acts as a transcriptional regulator, reducing the expression of VEGFR-2 and VEGFR-3 while increasing VEGFR-1 expression (82), which functions as a decoy receptor to inhibit angiogenesis (83).

Our results demonstrated that serum-derived exosomes from RHD patients promote a pro-angiogenic shift in endothelial cells. HUVECs treated with RHD patient’s serum-derived exosomes, showed elevated gene expression of VEGF-A (2.7-fold), MMP2 (1.5-fold) and IL4 (15.17-fold), all of which support angiogenic processes. Concurrently, expression of RASA1 (0.6-fold) and endostatin (0.4-fold), two angiogenesis inhibitors, was suppressed. This dual modulation upregulation of pro-angiogenic factors and suppression of anti-angiogenic genes suggests that RHD patient’s serum-derived exosomes create a favorable environment for abnormal vascular remodeling. These findings provide mechanistic insights into endothelial cell activation in RHD and highlight the role of exosomes as potential drivers of this pathology. Previous research from our lab revealed significantly unregulated miR-150-5p levels in the serum-derived exosomes from RHD patients, as determined by next-generation sequencing (NGS) analysis (57). We found up-regulated miR-150-5p (34.29 ± 9.524-fold mean difference; p= 0.0007) expression in RHD patient’s serum-derived exosomes as compared to those from healthy individuals. The role of miR-150-5p in various conditions, such as cancers, cardiovascular diseases (84, 85), anti-inflammatory and anti-apoptotic roles (86), is well-documented. Using miRDB, we predicted MYB and TP53 as potential targets of miR-150-5p, both implicated in angiogenesis regulation. Prior evidence suggests that miR-150-5p down- regulates TP53 (87). TP53, is a tumor suppressor gene, that down-regulates VEGF-A expression (88), thus affecting angiogenesis (89). In our study TP53 (0.227-fold; p= 0.006) expression was notably reduced in HUVECs treated with RHD patient’s serum-derived exosomes, while MYB expression remain unchanged. To further validate the pro-angiogenic role of miR-150-5p, we transfected HUVECs with miR-150-5p mimics (for miR-150-5p overexpression) and anti-miR- 150-5p (for mir-150-5p suppression). HUVECs transfected with mimics exhibited pronounced elongation, stretching and endothelial tube formation, confirming their angiogenic activation. In contrast, anti-miR-150-5p and suramin treated cells showed cell death. This result suggests that elevated miR-150-5p such as that seen in RHD patient serum-derived exosomes can directly drive angiogenesis, likely repression of TP53. The promotion of neovascularization in our *in-vitro* RHD model supports the hypothesis that miR-150-5p contributes to the vascular abnormalities observed in RHD patients.

In summary, our study provides evidences that exosomes derived from RHD patient’s serum promote angiogenesis through miRNA-mediated gene regulation, particularly involving miR-150-5p and its downstream effects on TP53. These findings not only advance our understanding of the molecular mechanisms driving RHD pathogenesis but also highlight potential therapeutic targets for modulating abnormal angiogenesis in affected individuals.

## 5. CONCLUSION AND FUTURE PRESPECTIVES

Our study provides compelling evidences that exosomes derived from the serum of RHD patients contribute to abnormal angiogenesis, a process that may play a key role in disease progression. We demonstrated that these exosomes promote endothelial cell activation, tube formation, and vascular network formation *in-vitro*, highlighting their pro-angiogenic potential. This effect appears to be mediated, at least in part, by the up-regulation of angiogenesis-related genes such as VEGF-A (2.7-fold), IL4 (15.17-fold), MMP2 (1.5 fold) and down-regulation of angiogenesis inhibitors including RASA1 (0.65-fold) and endostatin (0.4 fold). A particularly notable finding was the elevated expression of miR-150-5p (34.29±9.524-fold) in RHD patient’s serum-derived exosomes. The miR-150-5p appears to enhance angiogenesis by targeting TP53 (0.227-fold), a tumor suppressor gene known to regulate vascular growth. Functional assay using miR-150-5p mimics confirmed its role in promoting endothelial cell elongation, tube formation, and neovascularization, supporting the hypothesis that miR-150-5p might act as a key molecular driver in RHD-related vascular remodeling. These insights point toward miR-150-5p as a promising biomarker and potential therapeutic target for regulating angiogenesis in RHD. The use of exosome miRNAs as disease indicators also opens new avenues for non-invasive diagnostics and personalized therapeutic strategies.

Looking ahead, future studies should aim to include large cohorts of RHD patient to validate our current findings and enhance their clinical significance. Specifically, comparing miR-150-5p expression levels in serum-derived exosomes across different stages of RHD could provide valuable insights into its correlation with disease severity. Moreover, analyzing miR- 150-5p expression directly in RHD-affected mitral valves and comparing it with mitral valves obtained from cadavers could offer a deeper understanding of its tissue-specific role in disease pathology. In parallel, *in-vivo* models will be crucial for confirming the functional impact of miR-150-5p. Collectively, these efforts would advance the development of exosome-based biomarkers and novel therapeutic strategies aimed for early detection, improved management of RHD.

### Conflict of interest

The authors declare no conflicts of interest, either among themselves or with their affiliated institution.

## Acknowledgements

This study was conducted at the Department of Experimental Medicine & Biotechnology, PGIMER, Chandigarh, India, with financial support from the Council of Scientific and Industrial Research (CSIR), Ministry of Science and Technology, Government of India, under research grant no. 27(0352)/19/EMR-II.

## Notes

### Competing Interest Statement

The authors have declared no competing interest.

