## Supplementary Figures for "Serum exosomal miR-150-5p in rheumatic heart disease-associated angiogenesis": Supplementry.docx

**Supplementary File**

**Effect of RHD/CABG patient’s serum, PF and column chromatography purified exosomes on HUVECs**

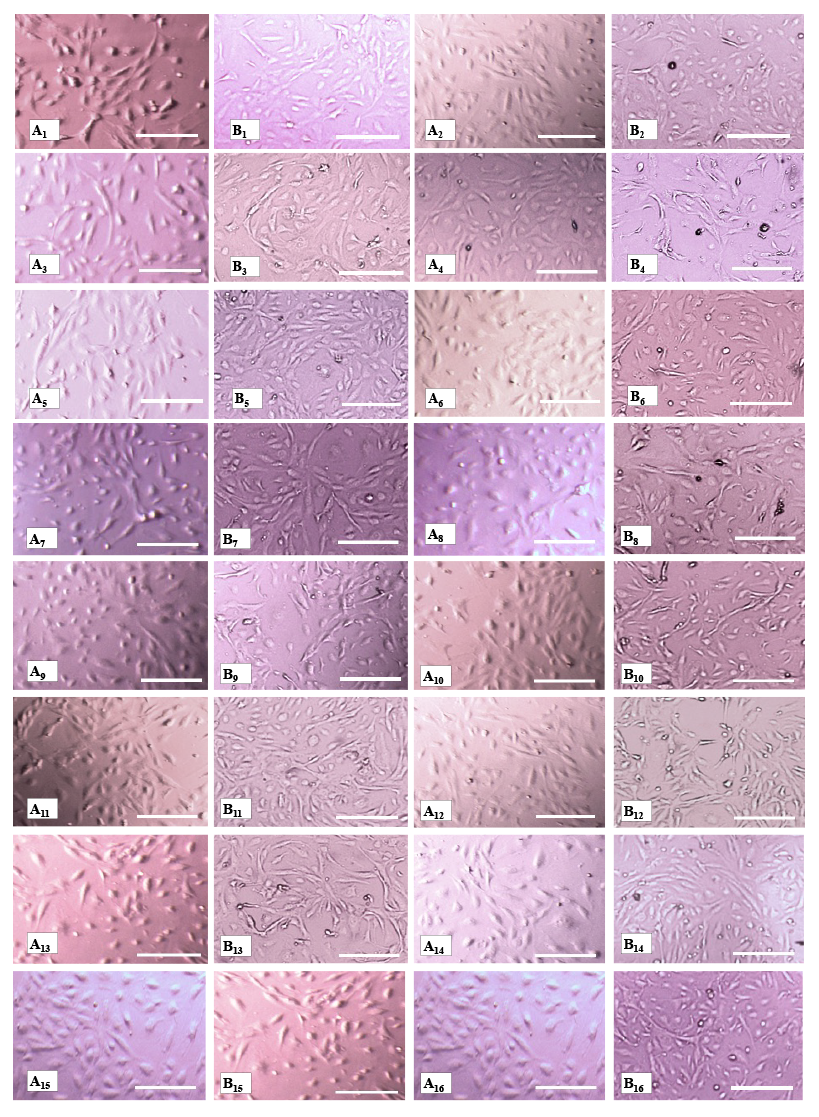

**Figure S.1: Angio-modulatory effects of exosomes on HUVECs:** (A) Morphology of HUVECs before treatment (0 hours) and (B) after 24-hour treatment with 100 µg of various serum and exosome preparations. HUVECs were treated with the following: (A_1_, B_1_) Placebo (10 mM potassium phosphate buffer, pH 8.0); (A_2_, B_2_) CABG pericardial fluid (PF); (A_3_, B_3_) CABG PF exosomes; (A_4_, B_4_) CABG PF + CABG PF exosomes; (A_5_, B_5_) CABG serum; (A_6_, B_6_) CABG serum exosomes; (A_7_, B_7_) CABG serum + CABG serum exosomes; (A_8_, B_8_) RHD PF; (A_9_, B_9_) RHD PF exosomes; (A_10_, B_10_) RHD PF + RHD PF exosomes; (A_11_, B_11_) RHD serum; (A_12_, B_12_) RHD serum exosomes; (A_13_, B_13_) RHD serum + RHD serum exosomes; (A_14_, B_14_) Healthy serum; (A_15_, B_15_) Healthy serum exosomes; (A_16_, B_16_) Healthy serum + healthy serum exosomes. All treatments were examined under an inverted microscope at 10X magnification (scale bar = 200 µm).

**Predicted target genes of miR-150-5p in HUVECs: MYB and TP53**

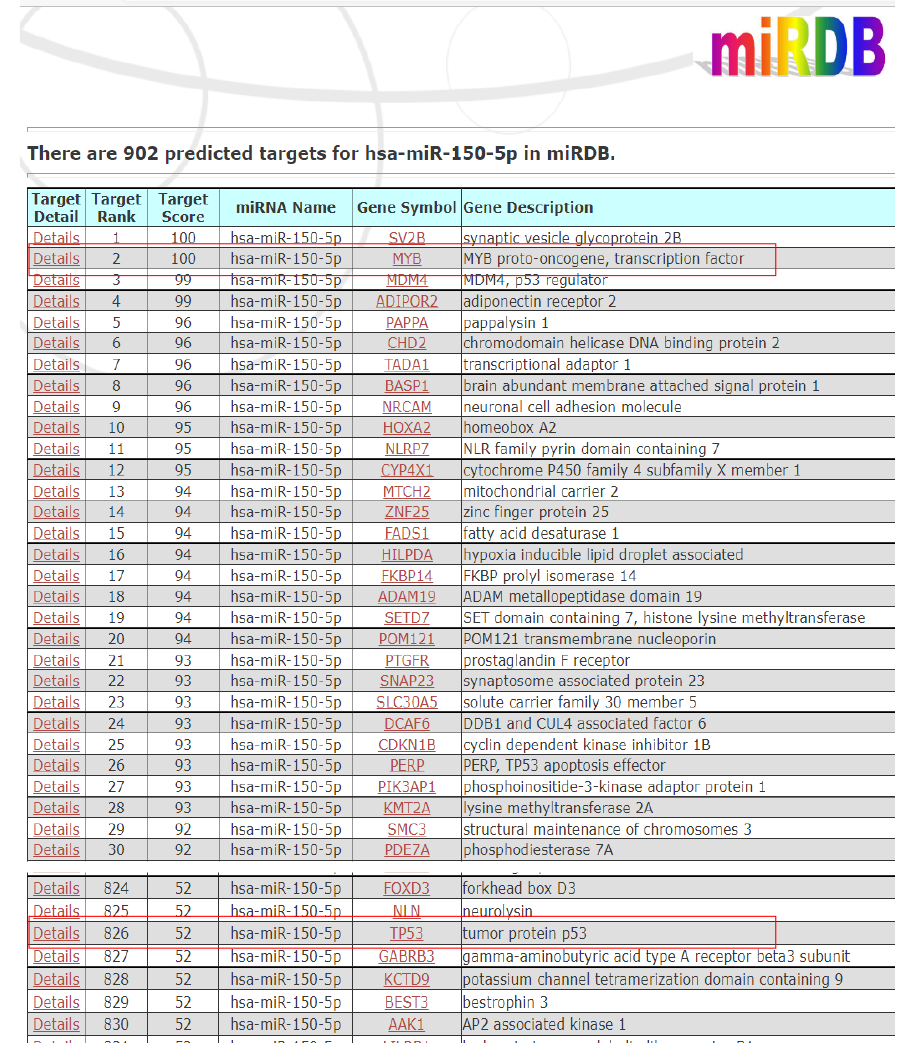

**Figure S.2: *In-silico* analysis using the miRDB database:** Identified **MYB** and **TP53** as potential target genes of miR-150-5p. This prediction supports the observed downregulation of TP53 gene expression in HUVECs treated with serum exosomes (upregulated expression of miR-150-5p) from RHD patients (**Fig. 5.B**).
